# Industrial Waste based Bio-manufacturing of Synthetic Tandem Repeat Protein Fibers

**DOI:** 10.1101/2020.03.16.994608

**Authors:** Tarek El-Sayed Mazeed, Huihun Jung, Yusuke Kikuchi, Benjamin D. Allen, David W. Wood, Melik C. Demirel

## Abstract

Protein-based fibers are lightweight, biodegradable, have excellent moisture and temperature regulation, and exceptional mechanical properties, but they are limited in production capacity. Biosynthetic protein-based fibers have the potential to overcome these concerns, but large-scale production with high yield (>1g/L) and purity (>%80), as well as low cost (<$50/kg), must be achieved. Here we developed an optimized expression and purification method for biosynthetic tandem repeat proteins, that are inspired from squid ring tooth (SRT) protein using three wet-waste feedstock, corn steep liquor, molasses, and soybean extract. SRT is composed of a highly stiff, naturally occurring bioplastic and these properties arise from the molecular architecture of the constituant proteins, which are segmented co-polymers with alternating semicrystalline and amorphous domains similar to silk. We have developed protocols to use liquid industrial and agricultural waste as feedstock for SRT production, which has the potential to divert waste streams into useful products. We also show that our biosynthetic protein powder, produced at 1 g/L yield and greater than 80% purity, can be manufactured into fibers using conventional split film or wet-spinning approaches.

## Introduction

Man-made synthethic fibers have been invented and commercially produced at large scale (350 million tons of plastics in 2017) in the last century (Geyer, Jambeck, & Law, 2017), but due to environmental concerns (e.g. toxic chemicals used in textile production, microfiber shedding that pollutes marine life, and contamination end-of-life disposal) there is a big demand for creating natural fibers from alternative sources (Jambeck et al., 2015). Since the dawn of civilization, natural fibers (e.g., wool, cotton, sisal, ramie, silk) have been used in textiles. However, due to increased population and cost issues, synthetic materials made of polyester, nylon, and others have replaced natural alternatives. (Drews, Barker, & Hatcher, 1985) The world-wide clothing textile market is extremely large, approaching $3 trillion. Many of the current processes are inexpensive, but their associated environmental costs can be enormous, and include petroleum consumption, carbon release, and microplastic pollution. (Hasanbeigi & Price, 2015) Hence, alternative production and processing technologies are needed in materials manufacturing (Zhu, Romain, & Williams, 2016). Natural fibers made by industrial biotechnology could give the textile industry and downstream markets the ability to operate more efficiently, cleanly, and profitably. (Moncrieff, 1966) To develop a successful, sustainable solution, the next generation of fibers should yield superior properties (e.g., mechanical, chemical, optical), durability and recyclability, reduce harm to nature, and address the environmental crisis with innovative, inspiring processes.(Pena-Francesch, Domeradzka, et al., 2018) A key requirement for the successful development of bioplastics-based fibers as a practical material is the ability to produce and purify them at large scale (e.g., ton scale) and at low cost (e.g., <$10/kg). (M. C. Demirel, Cetinkaya, Pena-Francesch, & Jung, 2015) Natural protein-based fibers are biodegradable (Shimao, 2001), but expression of recombinant proteins at industrial scale has proven challenging, particularly for structural proteins such as biosilk. (Peters, 1963) However, biosynthetic fibers have disadvantages due to low yields of production (e.g., due to high molecular weight), as well as complications related to the requirement of post-treatment (e.g. shear for recombinant silk) for solution processing in the production of fibers. (M. Demirel, 2019)

Peptide motifs from structural proteins such as silk(Dinjaski & Kaplan, 2016), elastin(Wise, Mithieux, & Weiss, 2009), collagen(Fratzl, 2008), keratin(Kayser, Grabmayr, Harasim, Herrmann, & Bausch, 2012), resilin(Elvin et al., 2005), and recently Squid Ring Teeth (SRT) protein (Jung et al., 2016a; V. Sariola et al., 2015), have been used to create multifunctional materials for diverse applications. Our focus has been on the study and modification of SRT in particular, for diverse applications in a variety of sectors. Various squid species have developed several features that allow them to be successful predators, which include sharp and rigid beaks, strong tentacles, and numerous SRT that allow these tentacles to grip prey. In particular, SRT are composed of a highly stiff, naturally occurring bioplastic with an elastic modulus (E) in the range of a 4-8 GPa and 100 MPa strength (Pena-Francesch et al., 2014). These properties arise from the molecular architecture of the constituant proteins, which are segmented co-polymers with alternating semicrystalline and amorphous domains.

Although the proteins that compose SRT tissues were only discovered recently, they have quickly gained the attention of several research groups due to their unique behavior. We have been exploring both native and recombinant SRT proteins and their biosynthetic variants in order to fabricate materials with tunable properties such as extensibility (Pena-Francesch, Jung, et al., 2018), biocompatibility (Leberfinger et al., 2017), thermal conductivity (Tomko et al., 2018), optical transparency (Yilmaz et al., 2017) and self-healing abilities (Veikko Sariola et al., 2015). SRT can be precisely tuned since a defined amino-acid sequence is genetically encoded in the DNA. This allows absolute control over stereochemistry, sequence, and chain length for tunable physical properties as well as environmentally friendly manufacturing. The mechanical durability and shedding resistance of SRT protein coatings are demonstrated in earlier studies by abrasion tests using a microfiber cloth as a model textile. These findings suggest that SRT coatings provide mechanical stability to textiles and could potentially prevent release of microfibers to the environment during mechanical abrasion.

Here we developed an optimized expression and purification method for biosynthetic tandem repeat proteins using industrial and agricultural waste as a feedstock. Further, we also show that our protein-based material, inspired from SRT sequences and produced via industrial fermentation, can be manufactured into fiberous forms using conventional split film or wet-spinning approaches. Biosynthetic fibers provide not only enviromentally sustainable production methods compared to synthetic fibers, but also opens up low-pollution methods of processing fibers for the future of textile industry.

## Results and Discussion

SRT is a high-strength thermoplastic protein that could be a green alternative to conventional plastics. However, in order to be industrially feasible, the expression levels of the SRT protein in well-established production hosts should be high (e.g., 1-10 g/L of fermentation volume), and it should be produced using an inexpensive medium (e.g., industrial waste). Further, the purification of the expressed SRT should also be simple and inexpensive, where chromatographic steps should be avoided if possible. Moreover, end-of-life options should be considered to determine the environmental impact of SRT, and the potential for enabling a circular economy of the product. In **Figure 1**, we conceptualize a circular-economy roadmap for SRT based on the use of industrial waste as feedstock, and the extent to which the final products (e.g. fibers) are reused or recycled by taking advantage of the reversible de-crosslinking of SRT proteins in organic solvents or weak acids.

**Figure 1:**
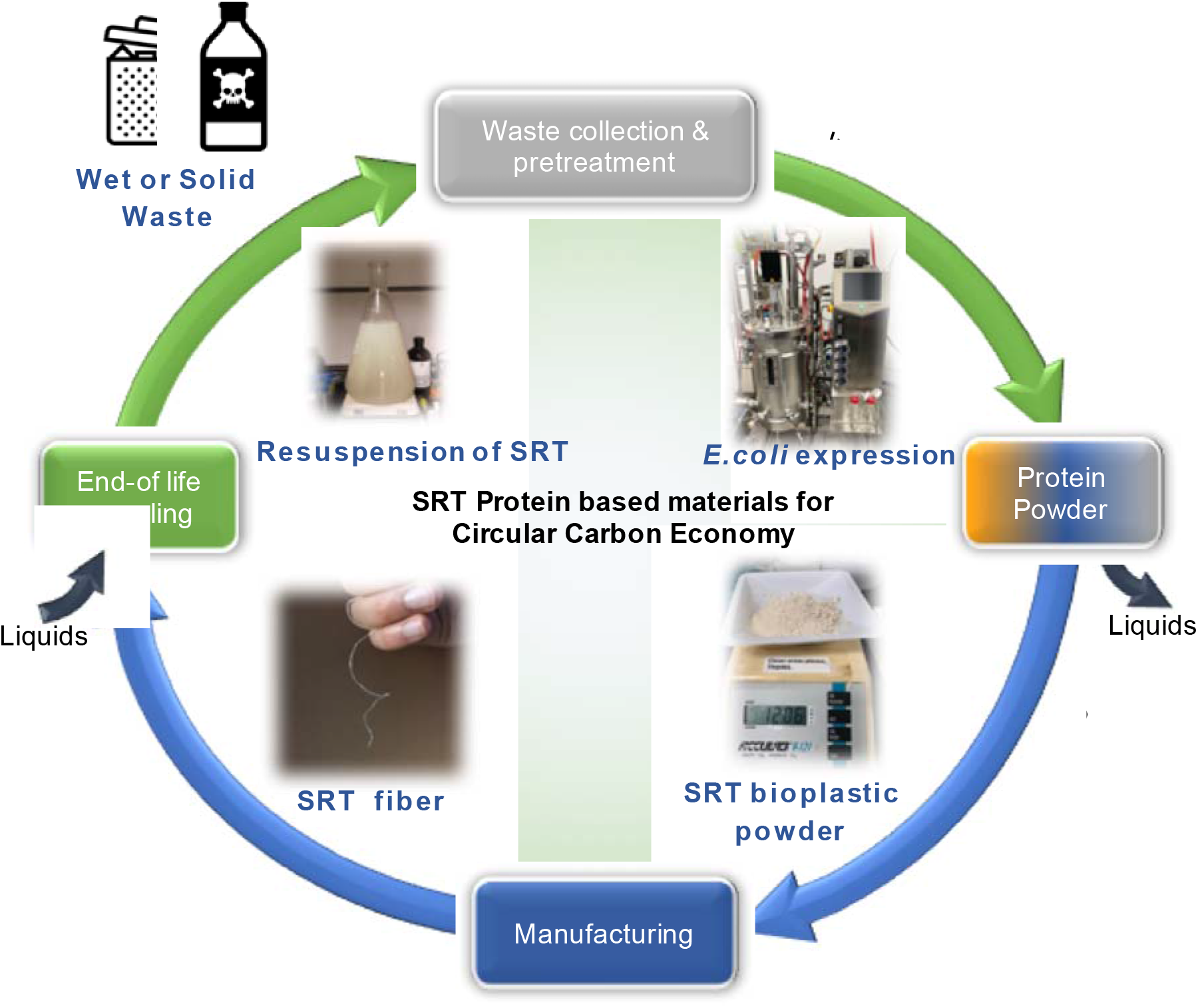
Circular carbon economy for squid ring tooth protein-inspired polypeptides produced from wet waste. SRT-based fibers are manufactured using fermentation using wet waste as feedstock and then purified using liquid separation. The protein powder is processed into fiber, which can be recycled via liquid processing.

Towards this circular-economy goal, we focused on the production of three sizes of the SRT tandem repeat (TR) protein, TR-n4, TR-n7, and TR-n11, referring to the sizes of the tandem repeat protein produced (molecular weights of 15, 25, and 42 kDa, respectively). Earlier, we developed a rolling-circle amplification method (Jung et al., 2016b) to produce tandem-repeat coding sequences in a single cloning step (**Figure 2a**). The designed building block for the tandem repeat proteins was based on the cross-linked crystal-forming sequence PAAASVSTVHHP and amorphous sequence YGYGGLYGGLYGGLGY observed in native SRT. We demonstrated earlier that the resulting polypeptides have similar semicrystalline β-sheet domains and amorphous tie-chains to the native squid proteins. As previously reported, the material properties of these proteins depend on their molecular weights, and we hypothesized that sequences of different lengths might require distinct conditions for optimal production.

**Figure 2:**
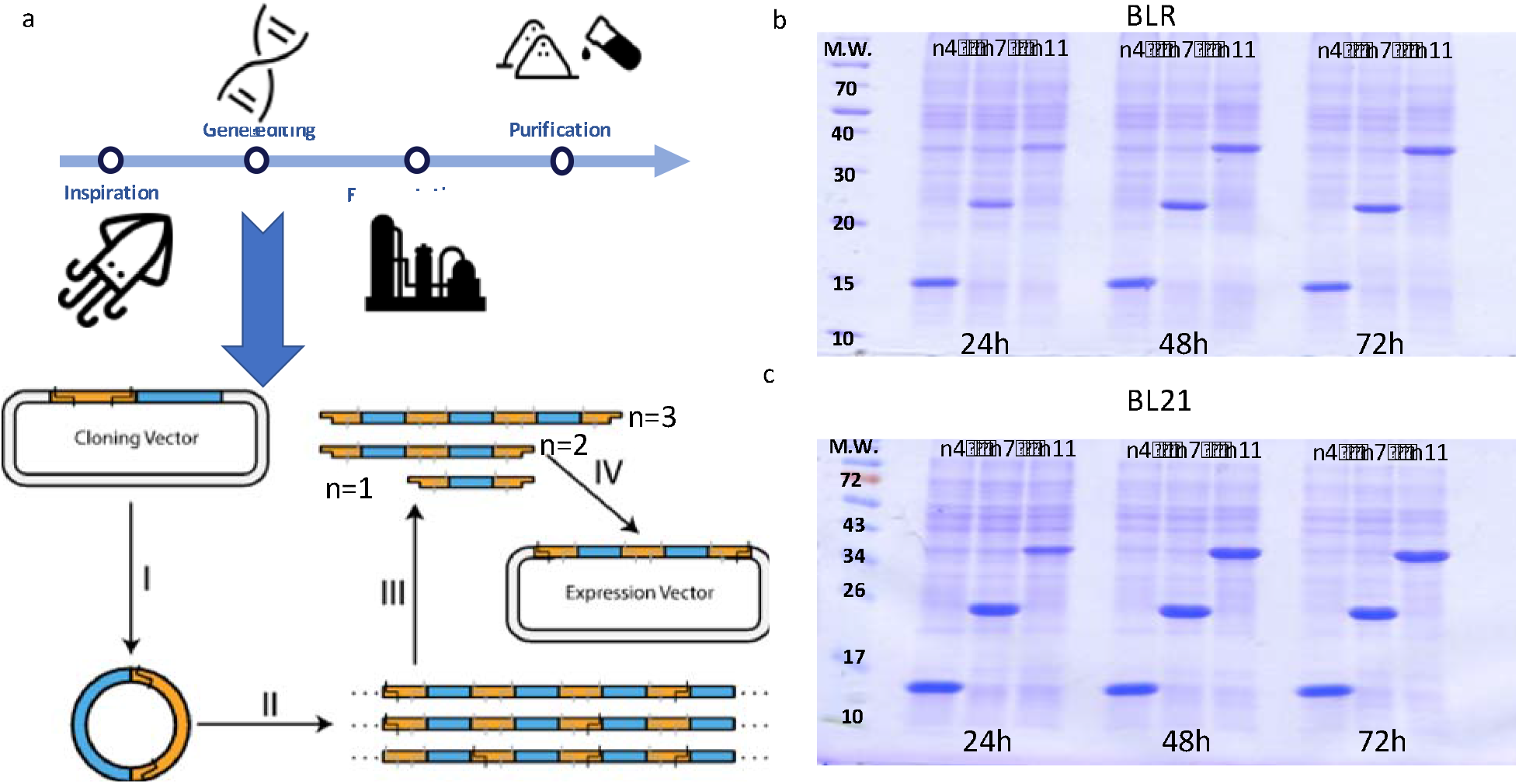
a) Overview of the design and preparation of tandem repeat polypeptides inspired by squid ring teeth. Rolling circle amplication is used to assemble repetitive genes. SDS-PAGE of cell-lysate samples taken at several time points for strains (a) BLR(DE3), and (b) BL21(DE3) bearing plasmid pET14b with inserts TR-n4, TR-n7, and TR-n11 incubated in 4X LB media at 37°C.

To produce these proteins, we worked with *E*. coli strain BL21(DE3) as well as BLR(DE3) as shown in **Figure 2b** and **2c** respectively. Because our protein-coding sequences are highly repetitive, we hypothesized that the recombination-deficient strain BLR(DE3) could enable more robust production. Since all of the proteins studied in this work are intrinsically insoluble under physiological conditions, our chosen production strategy was to allow slow accumulation of the TR proteins in inclusion bodies using the low-level uninduced expression provided by the pET14b plasmid in our chosen strains. The smallest of the proteins, TR15, shows early expression from the uninduced T7 promoter, and appears to reach its maximum production level by SDS-PAGE within 24 hours (**Figure S2**). The larger TR-n7 and TR-n11 proteins are expressed more slowly and appear to reach their maximum production levels by 48 hours of fermentation (**Figure 2b and 2c**). All three proteins remain stable during expression, with no loss of yield observed during 96 hours of incubation. Contrary to our expectations, strain BLR(DE3) produced uniformly less protein by SDS-PAGE across all conditions when compared to the standard strain BL21(DE3). This observation could be due to less efficient growth in the *recA*-deficient BLR strain, or perhaps protein production was affected by wider genomic differences between these strains that were not appreciated until recently. (Goffin & Dehottay, 2017)

It is known that transient anoxic conditions can cause limitations in amino acid production and plasmid stability. Hence we studied the effect of oxygen on TR-n11 production using a bioreactor with incubation at 37 °C and 500 rpm. The maximum production of TR-n11 was reached after 96 hours with agitation speeds of 500 rpm as shown in **Figure Supp-1**. The significant increase of TR-n11 expression by batch fermentation compared to the shake flask culture could explained by improved oxygen transfer in the bioreactor (Blibech et al., 2011). Qualitatively, the biomass in shake flask was lower than bioreactor production. (i.e., OD_600_ 21.7 in flask compared to OD_600_ 77.3 in bioreactor).

The adoption of SRT proteins for materials applications requires high-yield, high-purity production from inexpensive feedstocks. Hence, we screened alternative nitrogen sources using four (**Figure 3a**) selected agri-industrial residues at varying dilutions for their ability to serve as nutrients for TR-n11 production. Corn steep liquor (CSL) is a cheap potential source of nitrogen in fermentation, which is a major side-product in the corn starch industry. It is also a cheap source of proteins, amino acids, minerals, vitamins, and trace elements. Cane molasses is a by-product of sugar refineries, which contains saccharides and nitrogenous compounds, vitamins, and trace metal elements. The bagasse is constituted of lignin, hemicellulose, and cellulose. Three wet-waste sources, corn steep liquor (CSL), molasses, and soybean extract (SBE) enabled growth of our production strain and yielded TR-n11 protein, whereas growth in green juice extract was unsuccessful. **Figure 3b** and **3c** shows the protein gel (cell lysate and purified respectively) of four feedstock with varying dilutions. CSL feedstock growth medium reached the greatest production after 96 h of incubation.

**Figure 3:**
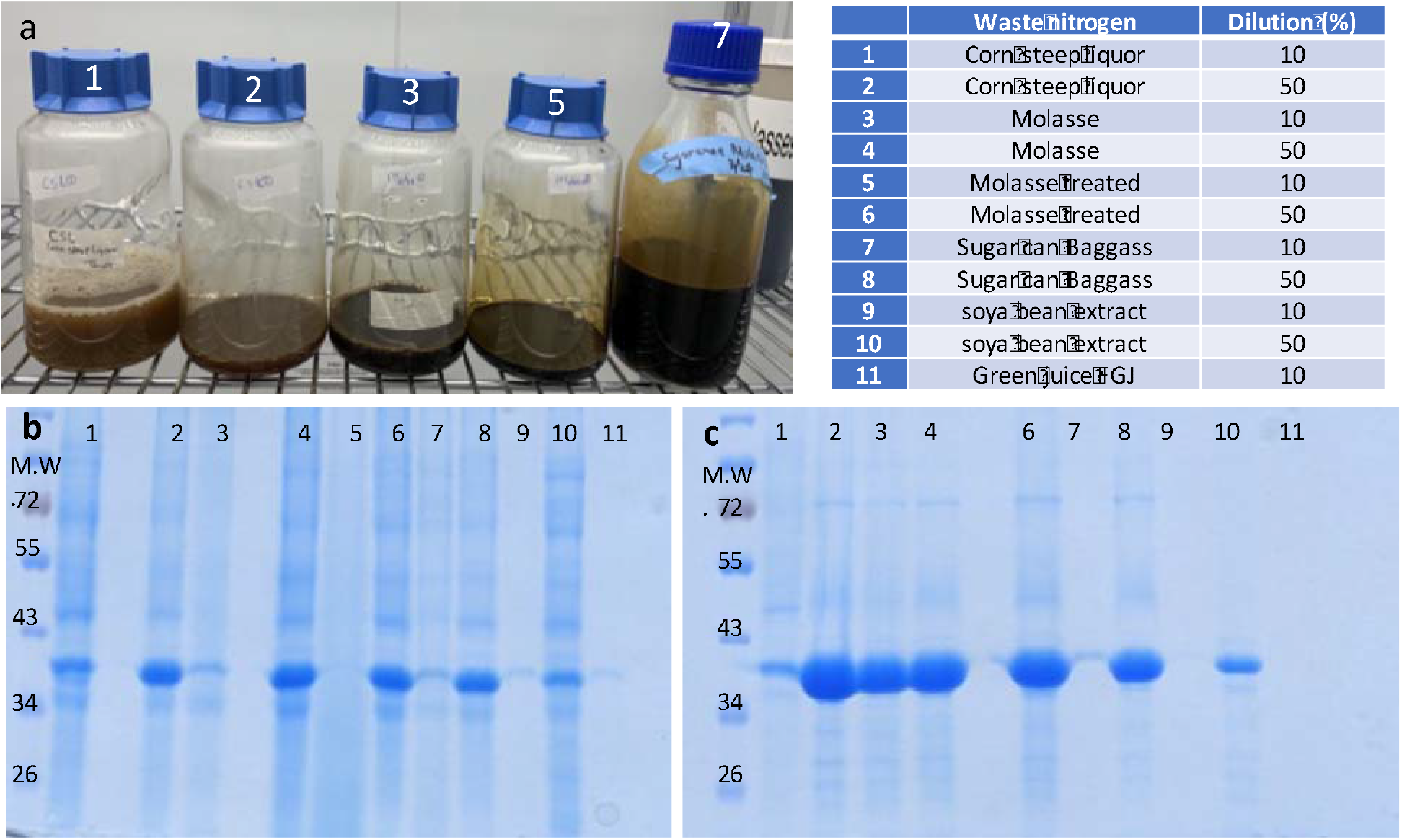
a) Pictures of corn steep liquor, mollase, sugar can baggass and soya been extract are shown. Cell-lysate (b) and buffer purified (c) samples of SDS-PAGE of TRn-11 (~40kDa) protein produced in BL21 (DE3) under un-optimized conditions from 11 different feedstock listed in the table.

Five assays were tested for production and purification of the recombinant TR-n11 protein as shown in Figure 4. Our initial approach to purify recombinant TR proteins was to simply wash bacterial inclusion bodies using a variety of detergents and solvents. Although this lead to modest increases in purity over cell lysate pellets, the purity was still less than 70% as judged by SDS-PAGE. We therefore attempted an organic extraction method where the inclusion body pellet was dissolved in DMSO and then re-precipitated using purified water as a counter solvent. This approach is based on the observation that TR proteins are highly soluble in pure DMSO, but become highly insoluble upon the addition of small amounts of water. Smaller protein domains, typical of the major contaminating proteins in the cell lysate, are expected to remain soluble in DMSO up to fairly high concentrations of water (up tp 200 mg/ml), which would afford a significant purification. As expected, this approach led to a substantial increase in purity, exceeding 70% (**Figure 4b**), which is also confirmed with Maldi (**Figure 4c**). The fact that this purity can be achieved in a single step using a highly scalable and simple method suggests that the unique properties of the SRT protein can be exploited to develop highly innovative and effective purification methods. We quantify production and purity of growth conditions using optical readouts of protein in 280nm, which is summarized in supplementary Table S1.

**Figure 4:**
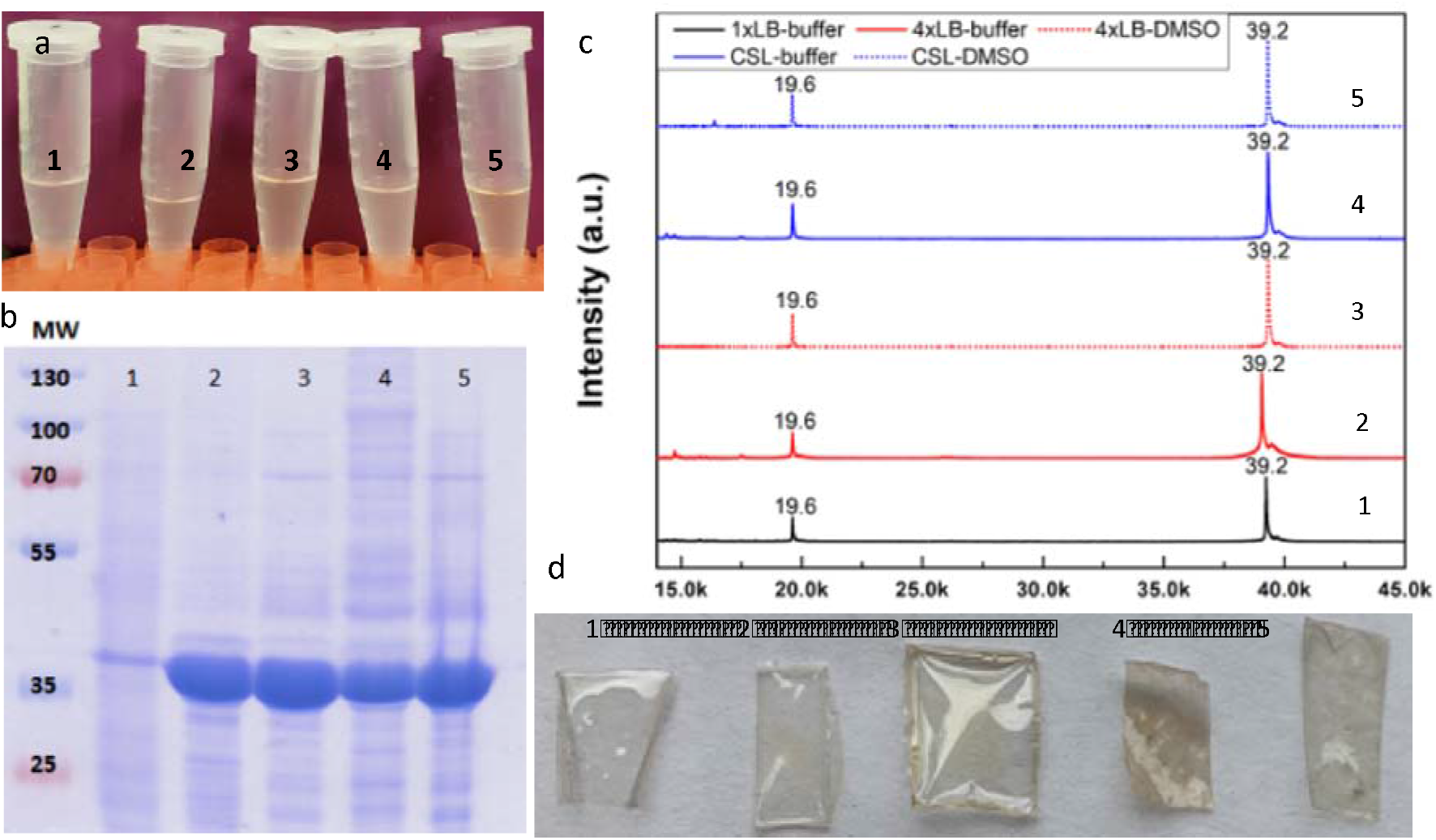
Five assays are tested for production and purification of the recombinant TR-n11 protein: 1) 1xLB feedstock and Buffer purification, 2) 4xLB feedstock and Buffer purification, 3) 4xLB feedstock and DMSO purification, 4) Corn syrup liquid (CSL) feedstock and buffer purification and 5) CSL feedstock and DMSO purification. a) All five production yield good solubility in HFIP solvent. SDS-PAGE of cell lysates shows high yield production of TRn-11 (~40kDa) for all except 1^st^ method. c) Maldi spectra also confirms the high purity of these samples. d) All five protein films were succesfully casted to form thin films that are mechanically robust.

Industrial waste based feedstock prodives unique advantage for producing inexpensive protein powder, which can be processed into fiber via solution processing. (Huang & Rha, 1974) In this study, we used two approaches, namely split film and wet spinning, to obtain protein fibers. Typically, proteins are exposed to various conditions (Traill, 1946) such as temperature, pressure, pH, and presence of solvents to process them from viscous solutions to solid fibers. (Lundgren, 1949) In split film fiber approach, TR-n11 is dissolved in hexafloroisopropanol (HFIP) as shown in **Figure 5a**. The protein solution is introduced into a saturated aqueous sodium chloride solution. The protein solution produced a thin film on the surface of the aqueous solution since SRT proteins are not water soluble. The protein film is twinned using a metal rod, and then washed and stretched until the material is capable of being converted into a fiber. In wet spinning (Paul, 1968), TR-n11 is dissolved in DMSO to convert the protein powder into a solution as shown in **Figure 5b**. The solvent is extruded through a needle injector by simply washing it out in aqueous bath. After extrusion, the solvent is removed and the filament is stretched to improve the pliability of the fiber. **Figure 5c** and **5d** show examples of TR-n11 fibers produced using split film fiber and wet spinning techniqes. TR-n11 are analyzed using FTIR in three conditions (i.e., unwashed, washed, and stretched) as shown in **Figure 5e and Figure S3a**. FTIR results revealed that these polypeptide chains contain ordered and amorphous domains. The amide I bands have been analyzed by using Fourier self-deconvolution and Gaussian fitting. FTIR peaks were assigned to secondary structure elements following the literature of fibrous proteins. The relative areas of the single bands were used in the calculation of the fraction of the secondary structure features. Each band is labeled as β-sheet (β), α-helix (α), random coil (rc), turn (t), or side chain (sc) according to the spectral regions of the amide I (1,600–1,700 cm^-1^) as shown in **Figure 5f and Figure S3b**. Analysis of the secondary structure content of the stretched TR-n11 film by Fourier self-deconvolution (FSD) and profile fitting revealed an increase in β-sheet content. Our earlier work of TR films suggests that high deformation induces chain alignment, β-sheet-crystal reorientation, and possible formation of ß-sheet fibrils within the protein matrix (Jung et al., 2016b).

**Figure 5.**
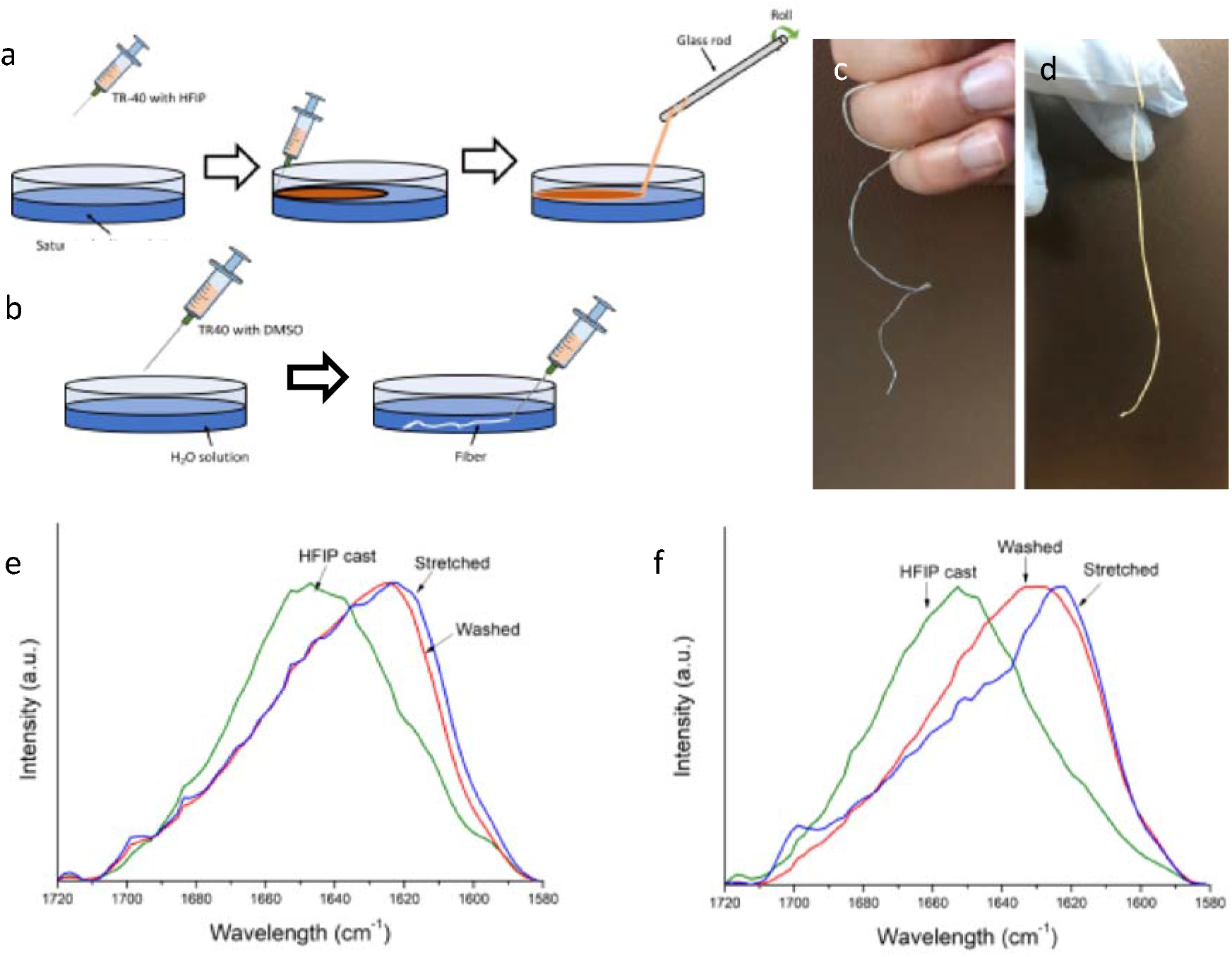
TR-n11 fibers and films are produced using solution processes: Schematic of (a) split film fiber, and (b) syringe based wet-spinning, and corresponding optical images of fibers are shown in c) and d) respectively. FTIR spectra of TR-n11 is investigated with e) hfip and f) dmso casted films under cast, washed and stretched conditions.

## Conclusion

In summary, we have developed eco-friendly processing and production technology based on industrial wet waste (i.e., corn steep liquor, molasses, and soybean extract) enabled growth that lowers the environmental impact of SRT based protein fiber production. Synthethic tandem repeat proteins inspired by SRT proteins are easier to produce, with 1 g/L yield and high purity (>%80), compared to other protein fibers due to their low molecular weights, also allow a variety of processing methods including split film and wet spinning processes, and offer novel physical properties compared to other natural and synthetic fibers such as self-healing and thermal switching. We envision sustainable textiles that are formed by protein-based fibers as the ideal choice due to their natural biodegradability, programmable physical and chemical properties, and reduced waste and energy demand compared to currently available fibers, if they can be scaled to industrial production using various waste sources (e.g., commercial, and residential sources or organic-rich wastewaters from industrial and commercial operations).

## Acknowledgments

The authors thank staff members of Penn State MRI and Huck User Facilities for helping data collection and sample characterization.

## Author contributions

M.C.D. and D.W conceived and designed the project. B.A. designed the tandem repeat constructs, H.J. and T.H. performed the protein expression and purification. T.H.. performed the industrial waste and optimization experiments, analyzed the data. Y. K. worked on fiber experiments. H.J. performed spectroscopic analysis of the protein materials. All authors participated in manuscript editing, revisions, discussions, and interpretation of the data.

## Data and materials availability

All data is available in the main text or the supplementary materials.

## Material and Method

### Chemicals and Reagents

All chemicals were purchased from either Sigma Aldrich (St. Louis, MO, USA) or Thermo Fisher Scientific (Waltham, MA, USA), unless otherwise stated. All cloning enzymes were purchased from New England Biolabs (Ipswich, MA, USA). All oligonucleotides were synthesized by Sigma Aldrich (St. Louis, MO, USA).

### Plasmid construction

TR proteins are constructed and expressed based on the protocols that are described earlier. (Jung et al., 2016b)

### TRs production with LB media

All protein expression experiments have performed in the *Escherichia coli* strains BLR(DE3) and BL21(DE3). For protein expression, strains transformed with expression plasmids (PET14 with TRn4, TRn7, or TRn11) were cultured in 5 mL Luria Broth (LB) media supplemented with 100 μg/mL ampicillin at 37 °C for 16 to 18 h. We used 3 different concentrations of LB media to produce TR proteien; 1 X LB (Yeast; 5g/l, Tryptone; 10 g/l, NaCl; 10 g/l), 2 X LB (Yeast; 10g/l, Tryptone; 20 g/l, NaCl; 10 g/l), and 4X LB (Yeast; 20g/l, Tryptone; 40g/l, NaCl; 10 g/l). The cultures were diluted 1:100 (v/v) into LB media different strength concentrations (1, 2, and 4X with keeping NaCl 1X). All protein (TRn4, n7, and n11) was dissolved in DMSO to a concentration of 50 mg/mL in a sonication bath for 1 h.

### Screening of different waste nitrogen source (agri-industrial wastes) for their use as nutrient in production media

Five different agri-industrial wastes, namely, corn steep liquor, sugar cane molasses, sugar cane bagasse, soybean extract and Green juice FGJ (Figure 3a) were evaluated as nutrients for TR42 production. All agri-industrial waste sources were obtained from Red Shield Acquisition, LLC. or Cargill Inc. All samples are used as feedstock after centrifugation and adjusting to pH to 7 after dilutions as stated in Figure 3a. The production media composed of 10%, 50% of the agricultural waste. Fermentation was carried out in 500mL shake flask containing 100 mL media at 37°C in an air shaker at 250 rpm for 120 h. Samples were taken at 24-h interval and TR42 content was examined using SDS-PAGE analysis of total protein.

### Sample preparation for SDS-PAGE with urea

Due to the intrinsic insolubility of the TR proteins studied in this work, a high concentration of the protein denaturant urea was required during sample preparation to observe the proteins by Coomassie-stained SDS-PAGE. Briefly, a 1 ml aliquot of bacterial culture or suspension of insoluble protein was centrifuged for 5 min at 13000 rpm and the supernatant was discarded. The pellet was then resuspended in 1 ml of aqueous urea (at 6, 9, or 15 M) and incubated at 90°C in a heat block for at least 10 minutes. To ensure that the sample was completely dissolved, the sample was vortex for 30 sec, and then 20 μl of solution was mixed with 20 μl 2x SDS-PAGE loading dye. The mixture was incubated for 5 min at 95°C, and 3 μl was used to to load a 10% SDS-PAGE gel.

### TR42 production in Bench-scale bioreactor

The bench scale was carried out in 14 l bioreactor (Eppendorf, New Brunswick Scientific Bioflo & celligen 115) with 9 l working volume and 37°C, 500 rpm, and aeration rate of 1.5 l min^-1^, as fermentation parameters. The pH was controlled at 7.0 with 40% (v/v) NaOH or H3PO4

### Tandem Repeat Protein purification

Three methods were developed for processing TR proteins.

#### (i) Dimethyl sulfoxide

First, harvest cells were resuspended in DMSO 3:1 with biomass by volume. Second, incubate in ambient conditions overnight and then spin down at 21200 g for 30 min at 25°C. Discard the precipitate and move the supernatant to a clean bottle, dilute the DMSO solution 1:1 with water by volume. Incubate in ambient conditions for 20 hr, and then spin down at maximum speed of 21,100 rcf for 30 min, collect precipitate, place it in-80°C overnight (or quickly freeze with liquid nitrogen) and place it in freeze drier for drying the sample(Labconco 7670520 FreeZone Legacy 2.5 Liter, USA) for 20 hr.

#### (ii) SDS purification

First, harvest cells were resuspended in in 1% SDS with 5:1 biomass by volume, and boil with stirrer for 30 min. Second, spin down and discard supernatant. After that resuspend the pellet with DI water and boil for 30 min. Repeat this washing step 3 times to make sure all SDS is removed from the pellet and stored at 80°C overnight. Move sample to freeze drier for 20 hr.

#### (iii) Buffer purification

This protocol has been adopted from Jung *et al.* (Jung et al., 2016b) Cell pellets were resuspended in 300 mL lysis buffer (50 mM Tris, pH 7.4, 200 mM, NaCl, 1 mM PMSF, and 2 mM EDTA) and lysed using a high-pressure homogenizer. The lysate was pelleted at 29,416 rcf for 1 h at 4 °C. The lysed pellet was washed twice with 100 mL urea extraction buffer [100 mM Tris, pH 7.4, 5 mM EDTA, 2 M urea, 2% (vol/vol) Triton X-100] and then washed with 100 mL washing buffer (100 mM Tris, pH 7.4, 5 mM EDTA). Protein collection in the washing step (urea extraction and final wash) was performed by centrifugation at 3,752 rcf for 15min. The resulting recombinant protein pellet was dried with a lyophilizer (FreeZone 6 Plus; Labconco) for 12 h.

